# Bacterial strain structure shapes the trajectory of antibiotic resistance genes from plasmid to chromosome

**DOI:** 10.64898/2026.04.13.718102

**Authors:** Martin Guillemet, Sonja Lehtinen

## Abstract

The evolutionary dynamics determining whether an antimicrobial resistance (AMR) gene resides on a plasmid or a chromosome are critical to understanding the spread of resistance. While theoretical models often predict that beneficial traits should ultimately integrate into the more stable chromosome, contemporary genomic data frequently shows a strong enrichment of resistance genes on mobile, yet costly and less stable plasmids. Here, we propose that the widespread plasmid-mediated resistance observed today may not represent a stable evolutionary equilibrium, but rather a snapshot of an ongoing process. Using a stochastic multi-strain model, we explore the role of strain diversity as a determinant of the timescale of this process. We suggest that the population structure of the bacterial host species, maintained by balancing selection, acts as a substantial barrier to the selective sweep of vertically transmitted chromosomal genes. Because horizontal transmission allows conjugative plasmids to readily cross these strain boundaries, our model indicates that plasmid-borne resistance can transiently dominate the population before chromosomal integration takes over, and that this transient dominance can last for decades. We illustrate these predictions by analysing antimicrobial resistance gene location in three common opportunistic gut pathogens. Furthermore, we show how extending the model to include co-selection between resistance genes on multi-drug resistance plasmids can lead to more complex dynamics that transiently reverse this plasmid-to-chromosome trajectory. Overall, our findings highlight how bacterial population structure can dictate the evolutionary trajectory of beneficial genes, suggesting that the current distribution of AMR genes between chromosome and plasmids is a prolonged transient state rather than a static endpoint.

## 1 Introduction

The evolutionary dynamics determining whether a bacterial gene resides on a plasmid or a chromosome are critical to understanding the spread of traits like antimicrobial resistance. Conjugative plasmids offer an advantage through their ability to move genes horizontally between lineages, facilitating rapid adaptation to new challenges [1]. However, this mobility typically comes at a cost: plasmids often impose a fitness burden on their host and are susceptible to segregation loss, making them inherently less stable vehicles for gene carriage than the chromosome [2, 3].

This fundamental trade-off has been the subject of extensive work. Empirical surveys have long established that gene location is non-random: ‘core’ housekeeping genes almost exclusively reside on the chromosome, while ‘accessory’ traits like antimicrobial resistance and virulence are frequently enriched on plasmids [4, 5]. Theoretically, several mechanisms have been proposed to explain why traits like antibiotic resistance are often plasmid-borne rather than chromosomal. Early models suggested that plasmids facilitate local adaptation, allowing genes to spread into immigrant lineages that lack the trait [6, 7]. Subsequent work has highlighted the role of fluctuating selection pressures [8], species diversity [9], or positive frequency-dependent selection, where the high horizontal transfer rate of plasmids allows them to establish a “priority effect” that prevents chromosomal integration [10]. However, a common feature of this literature is a focus on equilibrium states: identifying whether the gene is expected to reside on the plasmid or the chromosome in the long run.

Yet these equilibrium-based approaches may not capture the full picture of contemporary antimicrobial resistance. The widespread clinical and agricultural use of most antibiotics is a relatively recent selective pressure, primarily arising in the mid-20th century [11, 12]. The resulting increase in antibiotic resistance frequency has proved to be slow; for instance, large-scale genomic analyses have reconstructed the history of major resistance genes like *mcr-1*, revealing that they circulated for decades before becoming globally prevalent [13]. This suggests that the patterns of gene location observed in genomic data today may not represent a stable equilibrium, but rather a snapshot of a long and ongoing process [14, 15]. Understanding the nature and timescale of these transient dynamics is therefore critical.

Here, we study these transient dynamics and propose that intra-species host diversity plays a crucial role in the competition between plasmid- and chromosome-borne resistance genes. Most clinically important bacterial species are not monolithic clones but exist as diverse populations of coexisting strains [16, 17]. This strain structure is actively maintained by a variety of ecological and evolutionary forces [18], including resource competition [19], interference competition via bacteriocins [20], host-phage dynamics [21] or immunity [22, 23, 24]. While lower fitness costs and higher stability suggest that chromosomal resistance should be the favoured strategy at evolutionary equilibrium [3, 7] (although see [10]), we hypothesize that this population structure prevents or substantially delays the system from reaching this state. This delay arises from a fundamental asymmetry in transmission: while the high rate of horizontal gene transfer allows plasmids to easily cross strain boundaries [25], the population structure acts as a barrier to the population-wide selective sweep of a fitter chromosome-borne gene, which is largely limited to vertical transmission. Note that this framework applies specifically to acquired resistance genes and not to resistance arising from point mutations (e.g., *gyrA* mutations [26]).

To investigate this effect, we developed a stochastic multi-strain model. We focus on the scenario where i) resistance is first introduced on a plasmid (the most common route of introduction [12]), but is evolutionarily stable at equilibrium only on the chromosome, and ii) the plasmid is not maintained in the population as a pure parasite (i.e. without the resistance gene). We first estimate the impact of within-species diversity on the transient dynamics, with a particular focus on the time to fixation of chromosome-borne resistance. Then, we explore how a plausible extension to this model, i.e. considering multi-drug resistance, can alter these evolutionary trajectories. Finally, we compare these theoretical predictions to long-term evolutionary trends in public genomic data. Our work suggests that by slowing the displacement of transiently successful plasmids, population genetic structure is an important determinant of the non-equilibrium dynamics that characterize the contemporary evolution of antimicrobial resistance.

## 2 Model

We formulate the model using a set of ordinary differential equations (ODEs), represented in Figure 1, although we simulate the model stochastically using a tau-leaping approach (See Supplementary Note H). For each strain *i* ∈ {1, …, *n*}, we track the population density of four genotypes: sensitive cells (*S*_*i*_); cells carrying only the resistant plasmid (*P*_*i*_); cells with the resistance gene integrated into the chromosome (*C*_*i*_); and cells carrying both elements (*B*_*i*_).

**Figure 1:**
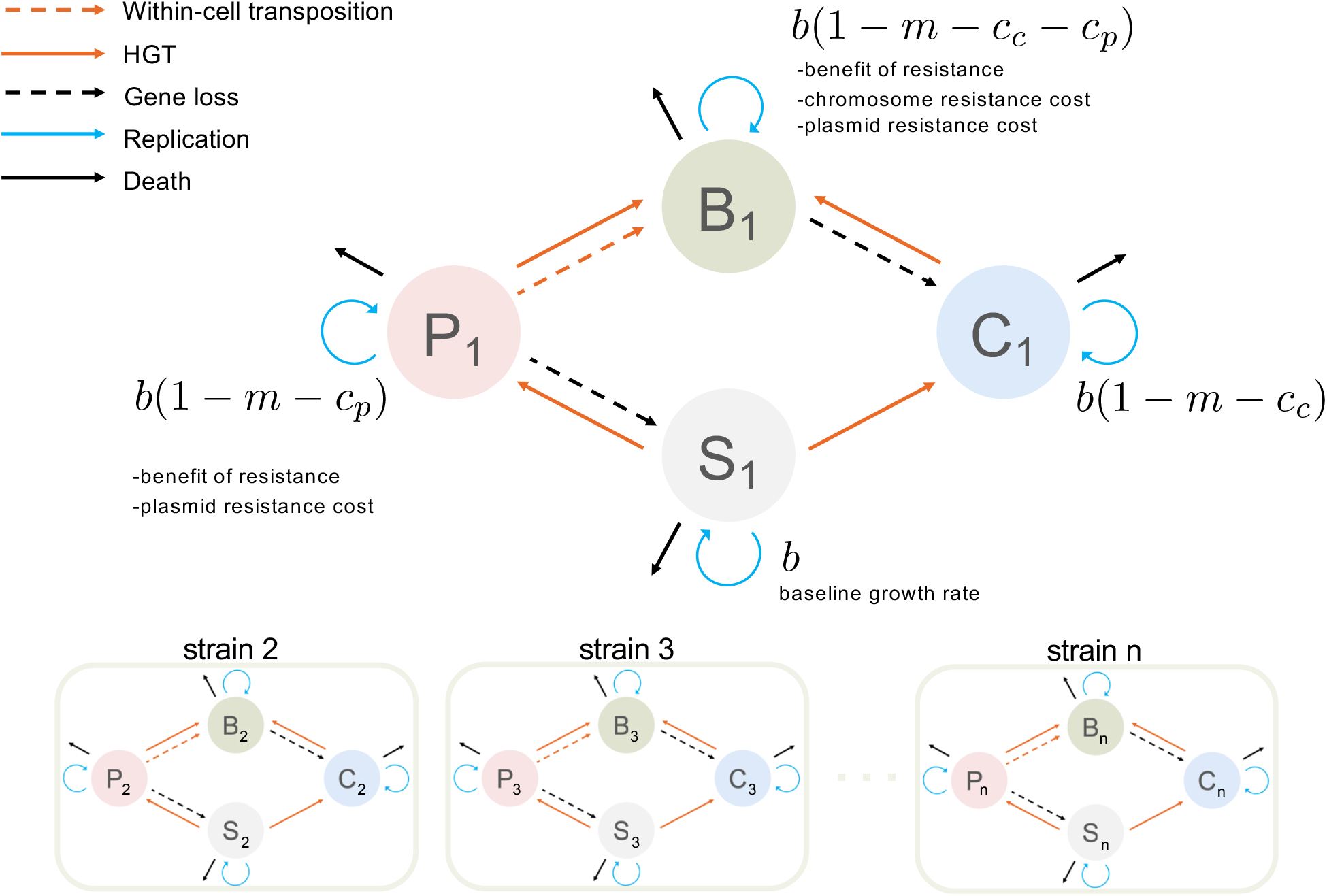
A multi-strain model of plasmidic and chromosomal resistance dynamics. The model tracks four genotypes within each of *n* competing strains: sensitive cells (*S*_*i*_, grey), cells carrying resistance on a plasmid (*P*_*i*_, red), the chromosome (*C*_*i*_, blue), or both (*B*_*i*_, green). Cells replicate (blue circular arrows) at a base rate *b*, modified by the benefit of resistance (*m*) and the costs of the plasmid (*c*_*p*_) and chromosome (*c*_*c*_) alleles. All cells are subject to a constant death/dilution rate (black arrows). The model includes several pathways for gene mobility: (i) horizontal gene transfer (HGT) (solid orange arrows), through plasmid conjugation or the less frequent between-cell chromosomal recombination; (ii) transposition (dashed orange arrows) which allows for intra-cellular “copy-paste” of the gene from the plasmid to the chromosome, creating a *B*_*i*_ cell; and (iii) gene loss (dashed black arrows) through the loss of the plasmid. This four-state dynamic is replicated across *n* coexisting strains, which are maintained in the population by balancing selection. Crucially, horizontal gene transfer is possible between strains, but the strain identity is only transmitted vertically.

The bacteria grow according to a logistic growth model with a base per capita division rate *b*, a shared carrying capacity *K*, and a fixed per capita death rate *d*. The per capita growth rate first depends on a strain-level balancing selection term, [1 − *η* (*f*_*i*_ − *f*_*eq*_)]. This produces balancing selection that pushes the frequency of each strain, *f*_*i*_, towards an equal equilibrium frequency, *f*_*eq*_. We denote

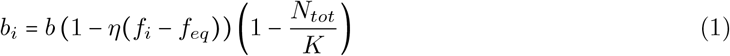

This captures the effects of balancing selection phenomenologically, whilst remaining agnostic about the specific mechanism giving rise to this selection. The division rate is further modified by the net benefit of resistance (*m*) and the costs of the elements carried (*c*_*p*_, *c*_*c*_). Cells with both plasmid and chromosomal resistance are subject to the cost of both elements, and the same benefit as cells with a single element.

The model incorporates several pathways for gene mobility. Resistance can spread horizontally between cells via plasmid conjugation (rate *γ*_*p*_) or chromosomal HGT (rate *γ*_*c*_). Additionally, the resistance gene can move between replicons within a single cell via “copy-paste” transposition. This is a replication-coupled event where a *P*_*i*_ cell can give rise to a *B*_*i*_ cell upon division with probability *q*_*T*_. Finally, the plasmid is subject to loss during replication, with probability *q*_*L*_. For simplicity, we neglect the loss of chromosome-borne resistance as the associated probability would be small [27].

The dynamics for any strain *i* are given by:

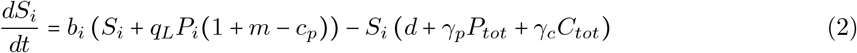

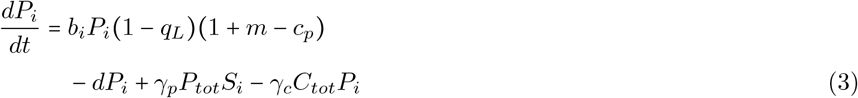

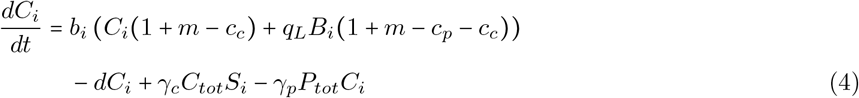

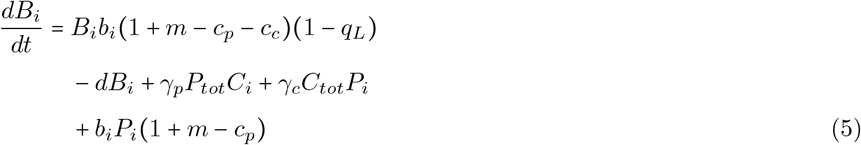

Here, 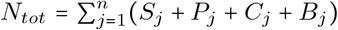 is the total population size. The total densities of plasmid-carriers (*P*_*tot*_) and chromosome-carriers (*C*_*tot*_) act as the donor pools for their respective HGT events.

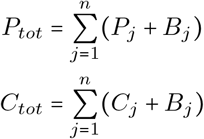

We parametrized the model using values derived from the experimental literature where available (Table 1). Although our model represents a simplified, chemostat-like system, the use of realistic parameters allows us to make semi-quantitative predictions regarding the timescales of the evolutionary processes under study. The absolute values for metrics such as fixation times presented throughout this work should therefore be interpreted not as precise predictions for complex natural systems, but as informed estimates of possible orders of magnitude and as internally consistent values for comparing different scenarios. We perform sensitivity analysis in the Supplementary Information (see Supplementary Notes C–F) for most parameters with a few exceptions. For instance, the impact of the fitness benefit of resistance *m* always involves the difference between this benefit and a cost *c*_*c*_ or *c*_*p*_, we chose to explore the sensitivity of the results to these cost parameters instead.

**Table 1:** Model Parameters and Default Values.

| Parameter | Description | Default Value | Justification & Reference | Sensitivity analysis |
| --- | --- | --- | --- | --- |
| ***Growth & Death*** |  |  |  |  |
| $b$ | Baseline per capita division rate. | $1.0\text{h}^{-1}$ | [28] | — |
| $K$ | Carrying capacity of the environment. | $10^8\text{ml}^{-1}$ | Based on typical bacterial densities in gut environments [29]. | Fig. S4 |
| $d$ | Per capita death/dilution rate. | $0.15\text{ h}^{-1}$ | A typical dilution rate used in chemostat experiments [30]. | — |
| ***Ecological Interactions*** |  |  |  |  |
| $n$ | Number of competing bacterial strains. | 20 | | Fig. 3 |
| $\eta$ | Strength of balancing selection. | 10.0 | Arbitrary strong value to maintain diversity and avoid strain extinction | — |
| ***Fitness Effects*** |  |  |  |  |
| $c_p$ | Fitness cost of carrying the plasmid. | 0.1 | Represents a moderate metabolic cost, consistent with experimental measurements [31]. | Fig. S3 |
| $c_c$ | Fitness cost of the chromosomal element. | 0.02 | Assumed to be lower than the plasmid cost. | Fig. S1 |
| $m$ | Net fitness benefit of resistance. | 0.2 | Both plasmid and chromosomal resistance have a net positive fitness effect. | — |
| ***Gene Mobility*** |  |  |  |  |
| $\gamma_p$ | Plasmid conjugation rate constant. | $10^{-12}\text{ ml}\cdot\text{h}^{-1}$ | Empirically measured rate for *E. coli* in liquid culture [32]. | Fig. S3 |
| $\gamma_c$ | Chromosomal HGT rate constant. | $10^{-19}\text{ ml}\cdot\text{h}^{-1}$ | Assumed to be several orders of magnitude lower than plasmid conjugation. [25] | Fig. S2 |
| $q_t$ | Probability of transposition per replication event. | $10^{-7}$ | Based on experimentally measured rates of transposition [27]. | Fig. S1 |
| $q_L$ | Probability of plasmid loss per replication event. | $10^{-4}$ | Represents a moderately stable natural plasmid [33]. | Fig. S3 |

## 3 Results

### 3.1 Invasion dynamics and the location of resistance

We now use our model to investigate the dynamics of plasmid- and chromosome-borne resistance when introduced into a sensitive population. We assume resistance is first introduced on a plasmid, the genetic element with the higher HGT rate. Figure 2 shows the dynamics of the invasion of a sensitive population when plasmid-borne resistance is introduced only in a single strain. When the population is homogeneous (Figure 2a,c), meaning that there is no maintained diversity through balancing selection, we find that the resistant plasmid quickly goes to fixation in the population (the analytical probability of successful plasmid invasion is derived in Supplementary Note A). Then, as soon as the resistance gene successfully transposes onto the chromosome and a resulting *B* cell loses the plasmid, the chromosome-borne resistance quickly outcompetes the plasmid, driving it to extinction (we analytically derive the probability of this evolutionary rescue pathway succeeding in Supplementary Note B). At equilibrium, all cells have chromosomal resistance. Note that double-resistant cells, carrying the resistance both on a plasmid and the chromosome, are not visible in Figure 2: they are less fit than single resistant cells and are quickly removed from the population. They must however be present transiently as they are an intermediate step for chromosome-borne resistance to emerge (as HGT for the chromosome-borne gene is quantitatively negligible, see Figure S2 and Supplementary Note D).

**Figure 2:**
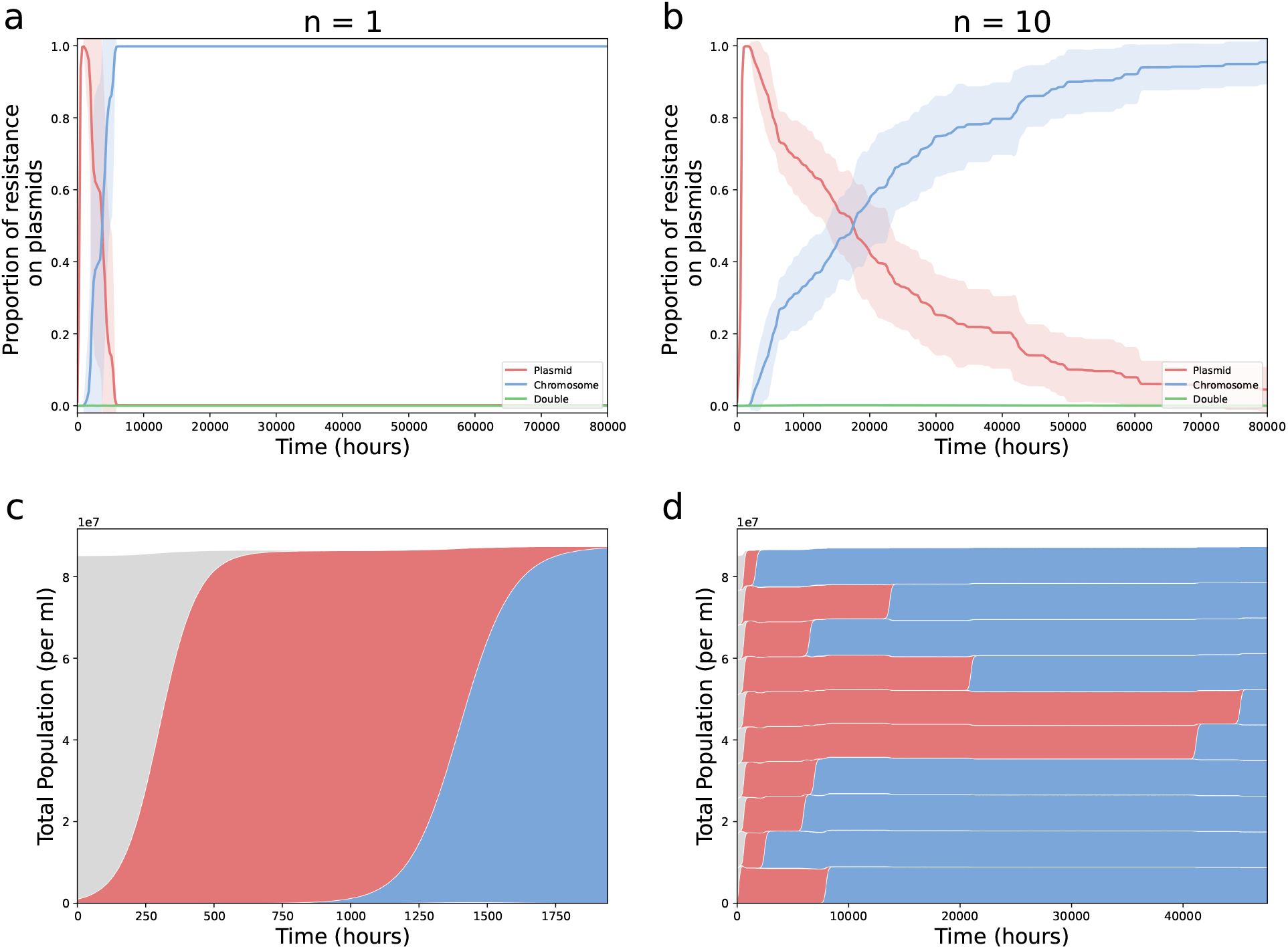
Evolutionary dynamics of plasmid-to-chromosome resistance transitions. Dynamics of resistance alleles in populations with single (*n =* 1) and high (*n =* 10) initial strain diversity. **(a-b)** Frequencies of plasmid-borne (red) and chromosomal (blue) resistance alleles over time, shown in **(a)** single-strain populations (*n =* 1) or **(b)** ten-strain populations (*n =* 10). Solid lines represent mean frequencies across 50 stochastic simulations and shaded regions denote ± SD. **(c-d)** Muller plots of individual simulation runs showing the succession of sensitive (grey), plasmid-resistant (red), chromosomal-resistant (blue), and double-resistant (green) lineages, shown for **(c)** a single-strain population (*n =* 1) or **(d)** across ten distinct strains (*n =* 10). All simulations are initialised with sensitive cells at their equilibrium density, alongside an inoculum of 10^6^ plasmid-resistant cells per ml. In the case of a structured population (**b**,**d**), this inoculum is done within a single strain.

When strain diversity is maintained in the population, the qualitative evolutionary trajectory remains the same, but the timescale of the dynamics is extended (Figure 2b). As in the homogeneous case, the plasmid initially invades and dominates due to its high mobility, but is eventually outcompeted and replaced by the fitter, more stable chromosome-borne resistance, which then goes to fixation. However, a major difference emerges in the timescale of this transition. For the chromosome to reach 99% frequency across the population, it takes approximately 13.2 times longer in the ten-strain population compared to the single-strain population (from 5, 000*h*± 1, 500 to 66, 000*h*± 25, 000, average over 10 simulations ± standard deviation). This extended timescale also prolongs the period of plasmid dominance, when the frequency of resistance is higher on the plasmid than the chromosome, by a factor 3.4 (from 4, 300*h*± 1, 400 to 14, 700*h*± 4, 000).

We can understand this delay by tracking the dynamics of each individual strain (Figure 2d). In the structured population, the displacement is not a single, population-wide sweep. Instead, it unfolds as a series of sequential, strain-by-strain invasions. The plasmid must first be introduced stochastically into each of the naive strains, after which the process of chromosomal emergence must occur independently within each of these newly established plasmid-carrying lineages. Each step of this sequential colonization adds to the total time, explaining the significant delay observed at the population level.

We investigate this trend in Figure 3, where we monitor the time to invasion as a function of the number of strains. We find that an increased number of strains increases the time until chromosome-borne resistance fixation (20-fold increase from 5 to 50 strains, *t*_5_ = 28, 000*h*± 16, 000 and *t*_50_ = 560, 000*h*± 170, 000). This also means that the transient time period in which plasmid-borne resistance dominates is longer when a population is divided into more strains.

**Figure 3:**
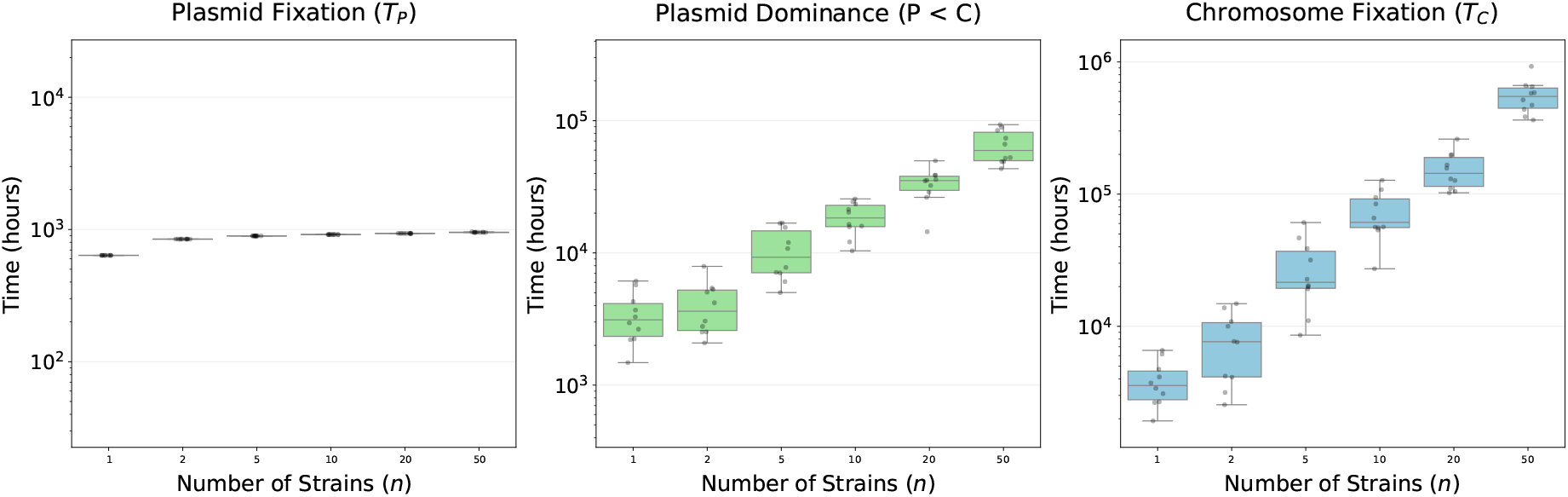
Strain diversity dictates the time to fixation of chromosome-borne resistance. Relationship between the number of strains maintained by balancing selection (*n*) and the time required for chromosomal resistance alleles to reach population fixation (using a frequency threshold of 0.99). Boxplots represent the median and interquartile range (IQR), with whiskers extending to 1.5× IQR. Overlaid circular markers indicate the results of individual stochastic simulation replicates (10 per condition). Simulations were initialized with a diverse sensitive population and a single strain with an inoculum of 10^6^ plasmid-bearing cells, with chromosomal resistance emerging via transposition. Fixation times are expressed in simulation time units (hours).

### 3.2 Linked selection on multi-drug resistance plasmids can drive chromosome-to-plasmid transitions

While our primary model shows a consistent trajectory from plasmid to chromosomal resistance, this need not necessarily be the case for more complex scenarios. In this section we explore one extension to the model: whether linked selection on a multi-drug resistance (MDR) plasmid could cause a gene already established on the chromosome to temporarily shift back towards plasmid-borne carriage. To test this, we simulated a population, similarly to the previous sections, where chromosomal resistance to Drug A (*C*_*A*_) goes to fixation. We then introduced a new MDR plasmid (*P*_*AB*_) carrying linked resistance to both Drug A and a second drug, Drug B (see Supplementary Note G for full model details).

The results demonstrate that linked selection can indeed drive a significant, albeit transient, increase in the proportion of resistance A residing on plasmids (Figure 4). At time zero, the frequency of Gene A on plasmids is near-zero. Upon introduction of the *P*_*AB*_ plasmid, this frequency rises sharply in both homogeneous (*n =* 1) and structured (*n =* 10) populations (Fig. 4a,b). During that phase, the MDR plasmid as a whole is beneficial because of the second resistance B, allowing it to increase in frequency. It is also mobile through conjugation, which allows it to spread between strains in the diverse *n =* 10 case. As this MDR plasmid spreads, it spreads the plasmid-borne copy of Gene A, causing the overall proportion of plasmid-borne Gene A to increase until it reaches 50%, where nearly all cells carry resistance A on both the chromosome and the MDR plasmid.

**Figure 4:**
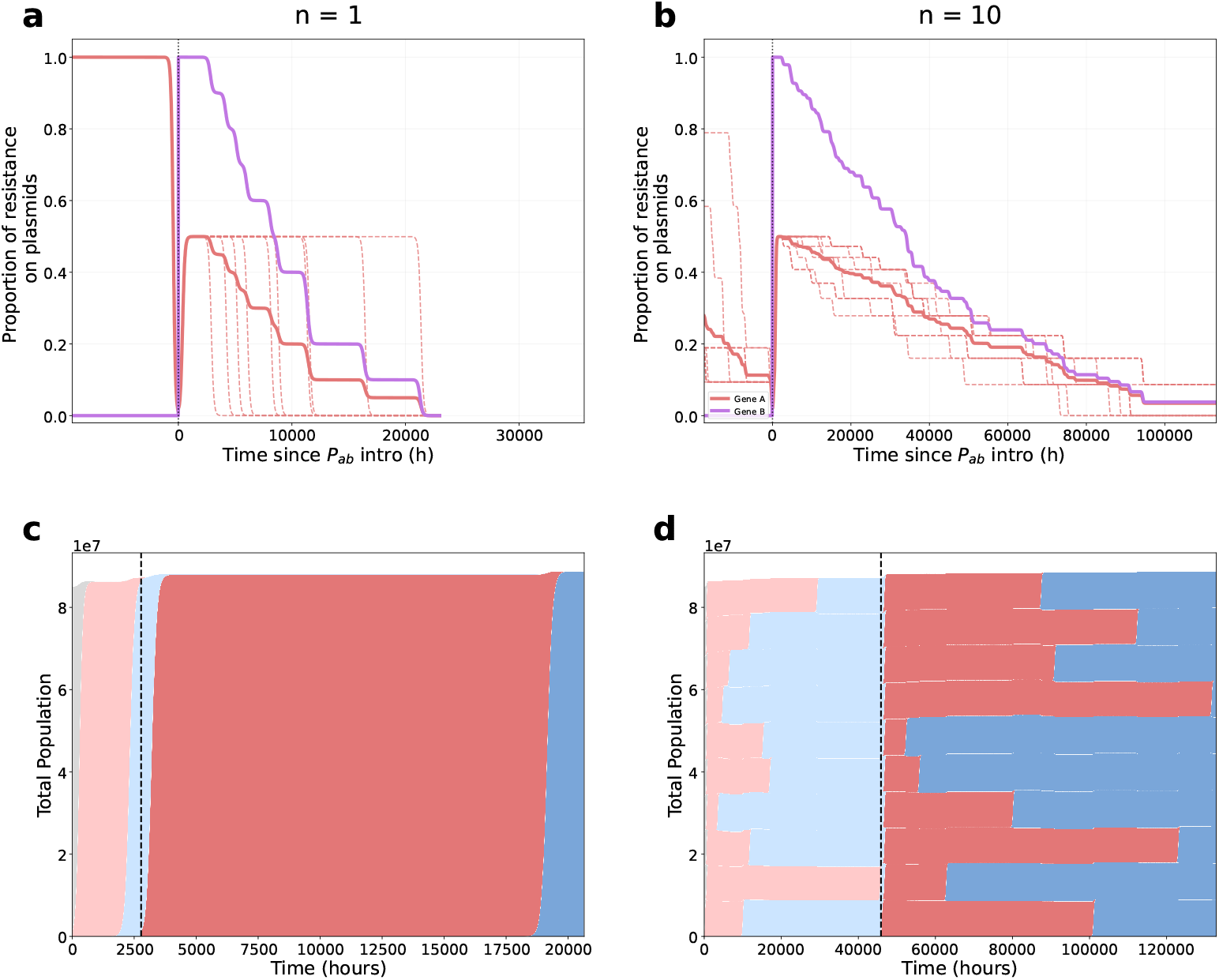
Evolutionary dynamics of the location of multi-drug resistance. Dynamics of resistance allele localization and lineage succession following the introduction of a multi-drug resistance (MDR) plasmid. **(a–b)** Proportion of resistance genes *A* (red, solid) and *B* (purple) residing on plasmids relative to the total resistant population. Time is aligned such that *t =* 0 corresponds to the triggered introduction of the *P*_*ab*_ plasmid (once chromosomal gene *A* frequency exceeds a 0.99 threshold). Thick lines represent mean values across 10 stochastic simulations; dashed red lines in the background indicate individual replicates for Gene *A*. Data are shown for **(a)** single-strain (*n =* 1) and **(b)** diverse (*n =* 10) populations. **(c–d)** Representative Muller plots illustrating the progression of genetic states within lineages. As in Figure 2, grey represents sensitive cell, light red and light blue respectively represent the cells carrying resistance *A* on the plasmid (*P*_*A*_) and chromosome (*C*_*A*_). Darker red represents cells with the multi-resistant plasmid and the *A* resistance on the chromosome (*P*_*AB*_*C*_*A*_), and darker blue corresponds to cells carrying both resistances on the chromosome without plasmid (*C*_*AB*_). Note that other types of cells are never in high enough density that they can be visible on the graph, and therefore are not associated with a color code. The vertical dashed line indicates the moment of *P*_*ab*_ introduction. Dynamics are shown for **(c)** *n =* 1 and **(d)** *n =* 10 initial strains. All simulations are initialised with sensitive cells at their equilibrium density, alongside an inoculum of 10^6^ plasmid-resistant *P*_*A*_ cells per ml. In the case of a structured population (**b**,**d**), this inoculum is done within a single strain. The *P*_*AB*_ is also introduced in the same quantity and in a single strain when it is triggered (when chromosomal gene *A* frequency exceeds 0.99).

Then, similarly to the one-resistance case, the B resistance gene transposes to the chromosome and the MDR plasmid is eventually lost within every strain. This effect is again amplified by population structure, as it forces the plasmid to chromosome transition within each individual strain, rather than once for the whole population in the *n =* 1 case. Overall, this modelled scenario shows how a plausible extension to our base model can alter the expected dynamics of resistance genes and produce more complex evolutionary trajectories.

### 3.3 Empirical evidence of chromosomal capture dynamics

Our theoretical model predicts that plasmids could serve as early spreaders for resistance invasion in strain-structured populations, but that they would eventually be outcompeted by chromosomeborne genes. A key semi-quantitative prediction of our stochastic simulations is the timescale of this transition: due to the hindering effect of strain structure, the replacement of plasmid-borne resistance by chromosomal integration is a slow process, estimated at the order of magnitude of 10^5^ hours (approx. 10 years) for moderate strain diversity (Fig. 3). This suggests that historical genomic data spanning the last decades would capture resistance genes in various phases of this slow evolutionary transition, rather than at a static equilibrium.

To test this, we analyzed the evolutionary trajectories of antimicrobial resistance (AMR) genes in *Escherichia coli, Klebsiella pneumoniae*, and *Salmonella enterica* using high-quality complete genomes curated from the NCBI MicroBIGG-E database. We applied binomial logistic regression to quantify the annual change in the log-odds of resistance genes being on plasmids.

The median regression coefficient is negative across all three species (*β*_med_ < 0, dashed blue lines in Fig. 5), indicating that for the average resistance gene, the probability of plasmid carriage has decreased over the last few decades. However, examining genes with statistically significant trends (Wald test, *P* < 0.05) reveals species-specific heterogeneity. In *Salmonella enterica*, all 13 significant genes exhibited a shift towards the chromosome compared to zero shifting towards plasmids (*n*_pos_ = 0). *Escherichia coli* displays an intermediate profile: while the overall trend indicates a shift towards the chromosome (*n*_neg_ = 9 vs *n*_pos_ = 5), a subset of genes are increasingly found on plasmids.

**Figure 5:**
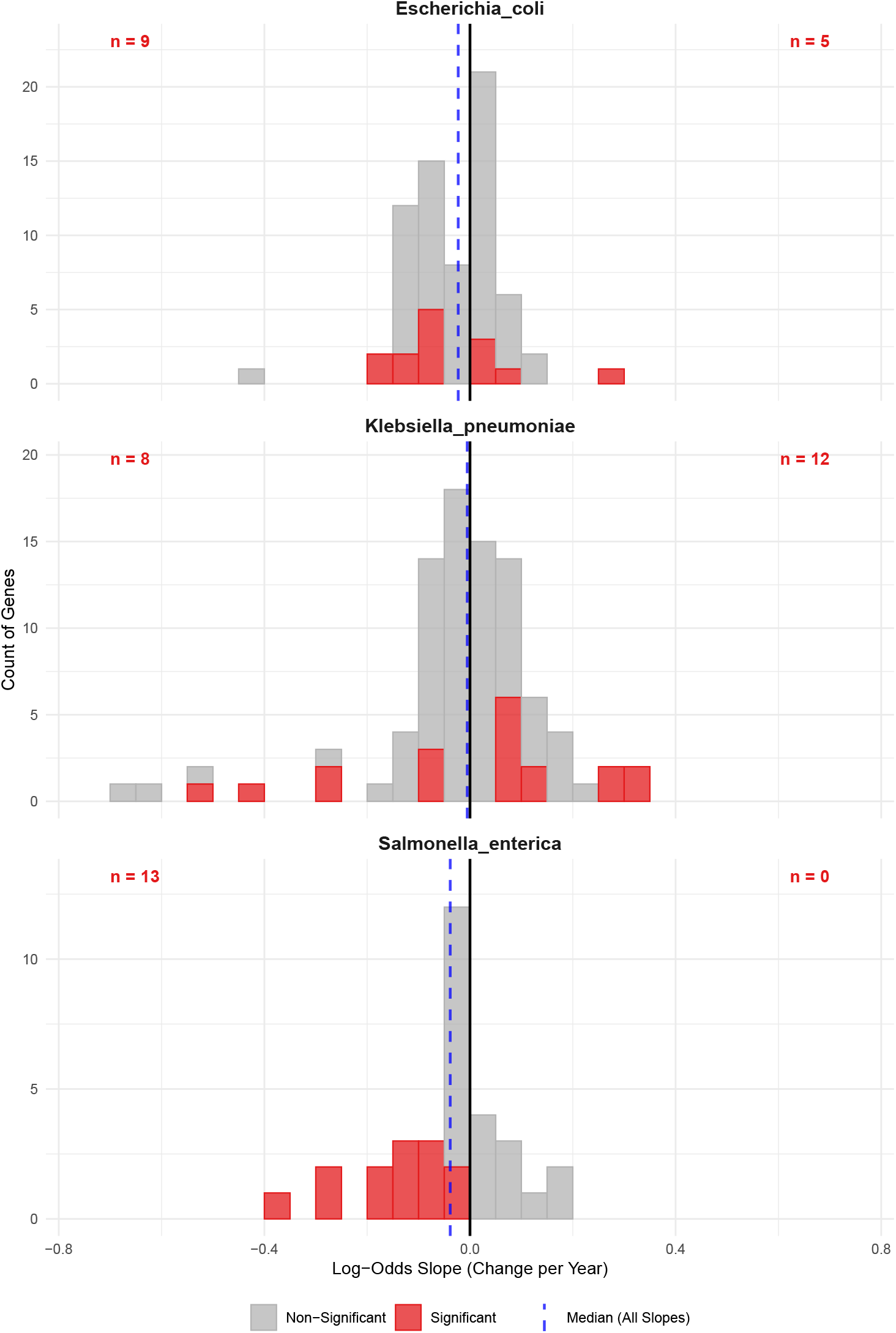
Temporal dynamics of plasmid-mediated resistance partitioning. Distribution of evolutionary trajectories for antimicrobial resistance (AMR) genes in *Escherichia coli, Klebsiella pneumoniae*, and *Salmonella enterica*. Histograms display the annual change in the log-odds of plasmid association (*b*_1_), estimated using binomial logistic regression on genomic data curated from NCBI (see Methods). Genes exhibiting statistically significant temporal trends (Wald test, *P* < 0.05) are highlighted in red; non-significant trends are shown in grey. Inset numbers (red) indicate the total count of genes exhibiting significant negative slopes (shift from plasmid to chromosome) or positive slopes (shift from chromosome to plasmid). Vertical reference lines indicate evolutionary stability (solid black line, *b*_1_ = 0) and the median slope for the species (dashed blue line).

In contrast, *Klebsiella pneumoniae* exhibits a distinct behavior. Despite a slightly negative median slope overall, the subset of genes with significant trends is dominated by shifts towards increased plasmid association (*n*_pos_ = 12 vs *n*_neg_ = 8). Within the framework of our model, this suggests that certain resistance genes in *E. coli* and *K. pneumoniae* (Figure 2b, early timepoints) could be evolving under more complex evolutionary pressures, for instance influenced by linkage with other resistance genes.

The magnitude of the observed slopes further supports the timescales predicted by our theory. Most statistically significant slopes fall within the range of |*β*_1_| ≈0.1–0.5. This corresponds to the probability of plasmid association dropping from 50% to 47.5-37.8% in one year. This gradual rate of change confirms that the displacement of plasmids by chromosomal resistance is not an instantaneous selective sweep, but a slow, decade-long process potentially constrained by population structure.

## Discussion

In this study, we combined stochastic multi-strain modeling with an analysis of longitudinal genomic data to investigate the evolutionary trajectory of antibiotic resistance genes. While chromosomal integration could be the long-term evolutionary optimum for many resistance genes, our central finding is that intra-species strain diversity acts as a powerful brake on this process. By forcing chromosomal genes to emerge and sweep independently within competing lineages, population structure extends a transient phase of plasmid dominance from negligible timescales to periods spanning potentially decades. Consequently, the plasmid-mediated resistance observed in contemporary genomic data may not represent a stable equilibrium, but rather a snapshot of a prolonged, non-equilibrium transition. Our empirical analysis of genomic data provides general support for this prediction, consistent with previous studies of beneficial gene location trajectories [14, 15]. The median evolutionary trend across *E. coli, S. enterica*, and *K. pneumoniae* indicates an overall gradual shift of resistance genes from plasmids towards the chromosome. The slow rate of this change confirms that the chromosomal capture of resistance is not an instantaneous selective sweep, but a slow and ongoing process spanning decades.

Previous theoretical and empirical work has identified several distinct evolutionary forces that dictate gene location. For example, the complexity hypothesis posits that core “housekeeping” genes involved in processes like replication and translation are part of large genetic networks and evolutionarily stuck on the chromosome, while other genes (like AMR) can be mobile [5, 34]. For mobile genes, the gene dosage hypothesis suggests that plasmids may be the optimal location for genes that benefit from high copy number, such as low-efficiency enzymes or efflux pumps [35]. Furthermore, while our model assumes a constant benefit to resistance, fluctuating selection can fundamentally alter these dynamics. Theoretical models suggest that if environmental pressures vary rapidly, the inherent segregational instability of plasmids can become advantageous, allowing populations to rapidly shed costly resistance genes when they are no longer needed [8]. Moreover, we consider a scenario without a competing sensitive plasmid, which has been shown to lead to priority effects where antibiotic genes tend to remain stable at equilibrium on the genetic element they were first introduced on [10]. Conversely, empirical studies have shown that compensatory evolution can rapidly ameliorate plasmid fitness costs, effectively stabilizing them in the population even in the absence of selection [36, 37]. Thus while the overall evolutionary forces impacting gene location are more complex than captured in our model, our results emphasize the relevance of non-equilibrium dynamics in thinking about gene location and the importance of strain structure in the timescale of these dynamics.

Despite the simplifications of our model, we propose that our central conclusions are robust. The core mechanism we highlight, that a diverse population of competing strains hinders the selective sweep of vertically transmitted alleles, must still be relevant in more complex settings, considering previous theory and experiments. Most of the real-world complexities not included in our model, such as spatial structure or intermittent selection, would likely not accelerate the fixation of chromosomal resistance and are more likely to slow it down even further. Therefore, the decade-long timescales predicted by our model are more likely conservative, lower-bound estimates for the duration of the transient plasmid-dominance phase in nature.

While the general trend in our empirical data aligns with our model’s long-term prediction of chromosomal capture, the data also reveal significant heterogeneity among species (Fig. 5). *Salmonella enterica* shows a strong and uniform trend towards chromosomal integration. In contrast, both *E. coli* and particularly *K. pneumoniae* display a more mixed profile, where a substantial number of resistance genes show a statistically significant, increasing association with plasmids. This observation of a reversed chromosome-to-plasmid trajectory appears to contradict the fundamental prediction that the chromosome is the more stable long-term location for a beneficial gene. As suggested, there are many mechanisms at play that could lead to that outcome. We explored one such plausible mechanism: the effect of linked selection on multi-drug resistance (MDR) plasmids. Our base model considers the dynamics of a single resistance benefit, but clinical environments often exert selective pressure from multiple antibiotics simultaneously [38]. Plasmids are uniquely capable of accumulating multiple resistance genes into single adaptive modules that provide a large, aggregated fitness benefit [1, 39]. We find that in a scenario where an MDR plasmid (*P*_*AB*_) is introduced in a population already carrying one resistance on the chromosome, we can observe a transient trend where the frequency of resistance shifts from chromosome to plasmid.

The empirical analysis we present serves as a proof-of-concept to illustrate that these long-term trends can indeed be visible in public genomic data. To this end, we employed highly stringent criteria for classifying gene locations, ensuring a high-confidence dataset, albeit at the cost of size. We also restrict the analysis to only three bacterial species: all common, opportunistic pathogens, and colonizers of the human gut. While a full investigation is beyond the scope of this work, our exploratory simulations of multi-drug resistance (MDR) plasmids suggest that linked selection could provide a plausible mechanistic explanation for some gene trajectories.

Throughout this study, we have framed our model and interpreted our results through the lens of antimicrobial resistance. We chose this focus for several reasons: AMR is a pressing global health crisis, its evolution is frequently mediated by plasmids, and the wealth of genomic data associated with it provides an opportunity to test theoretical predictions against real-world evolutionary trajectories. However, the fundamental mechanisms we model are not specific to antibiotic resistance. The evolutionary trade-off between a mobile, costly plasmid and a stable chromosomal allele applies to any recently introduced beneficial gene. While AMR provides a relevant case study, we hypothesize that the role of population structure in prolonging transient plasmid dominance represents a general dynamic in microbial evolution that governs the fate of a wide range of adaptive genes carried on mobile genetic elements.

## Methods

### Data Acquisition and Curation

Longitudinal data on the genomic distribution of antimicrobial resistance (AMR) genes were sourced from the NCBI Pathogen Detection MicroBIGG-E database (accessed December 2025). The analysis focused on three high-priority pathogens typically found as commensals in the gut microbiome: *Escherichia coli, Klebsiella pneumoniae* and *Salmonella enterica*.

AMR entries were cross-referenced against the NCBI RefSeq Assembly catalog. Only isolates belonging to assemblies designated as “Complete Genomes” were retained.

### Replicon sorting

The replicon type (plasmid vs. chromosome) for each resistance gene was determined through a hybrid approach combining assembly metadata and text mining. For every contig hosting an AMR gene, the assembly statistics and metadata were retrieved via the NCBI Entrez API using the `rentrez` R package. Classification followed a hierarchical decision tree:

1. **Number of contigs per assembly:** Assemblies consisting of exactly one scaffold (*n*_scaffolds_ = 1) were automatically classified as chromosomal, as these represent fully closed genomes with no extrachromosomal elements.
2. **Contig definition:** For multi-scaffold assemblies, contigs were classified based on explicit keywords in their NCBI definition. Definitions containing the term “plasmid” were assigned to the plasmid partition. Definitions containing “chromosome” or “complete genome” were assigned as chromosomal.

Contigs that could not be resolved by these criteria were excluded from the analysis.

### Statistical Analysis of Evolutionary Trends

To quantify the temporal dynamics of plasmid-mediated resistance, we modeled the genomic location of resistance genes using binomial logistic regression. This approach avoids the artifacts associated with linear regression on frequency aggregates by preserving the sample size weight of each observation year. Data were grouped by species, year, and gene. For each gene, the probability of being plasmid-borne was modeled as:

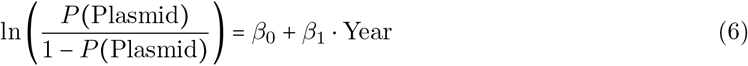

The slope coefficient *β*_1_ represents the annual change in the log-odds of the gene being located on a plasmid. Positive slopes indicate an evolutionary trajectory towards an increasing plasmid-borne frequency, while negative slopes indicate increasing chromosome-borne frequency. Genes with fewer than 100 total observations across the study period were excluded. Significance was assessed using Wald *Z*-tests, and 95% confidence intervals were calculated using the Wald method. All analyses were conducted in R (v4.4.0).

## Data availability

All relevant data and code are available at https://doi.org/10.5281/zenodo.19109071.

---

**Figure S1:**
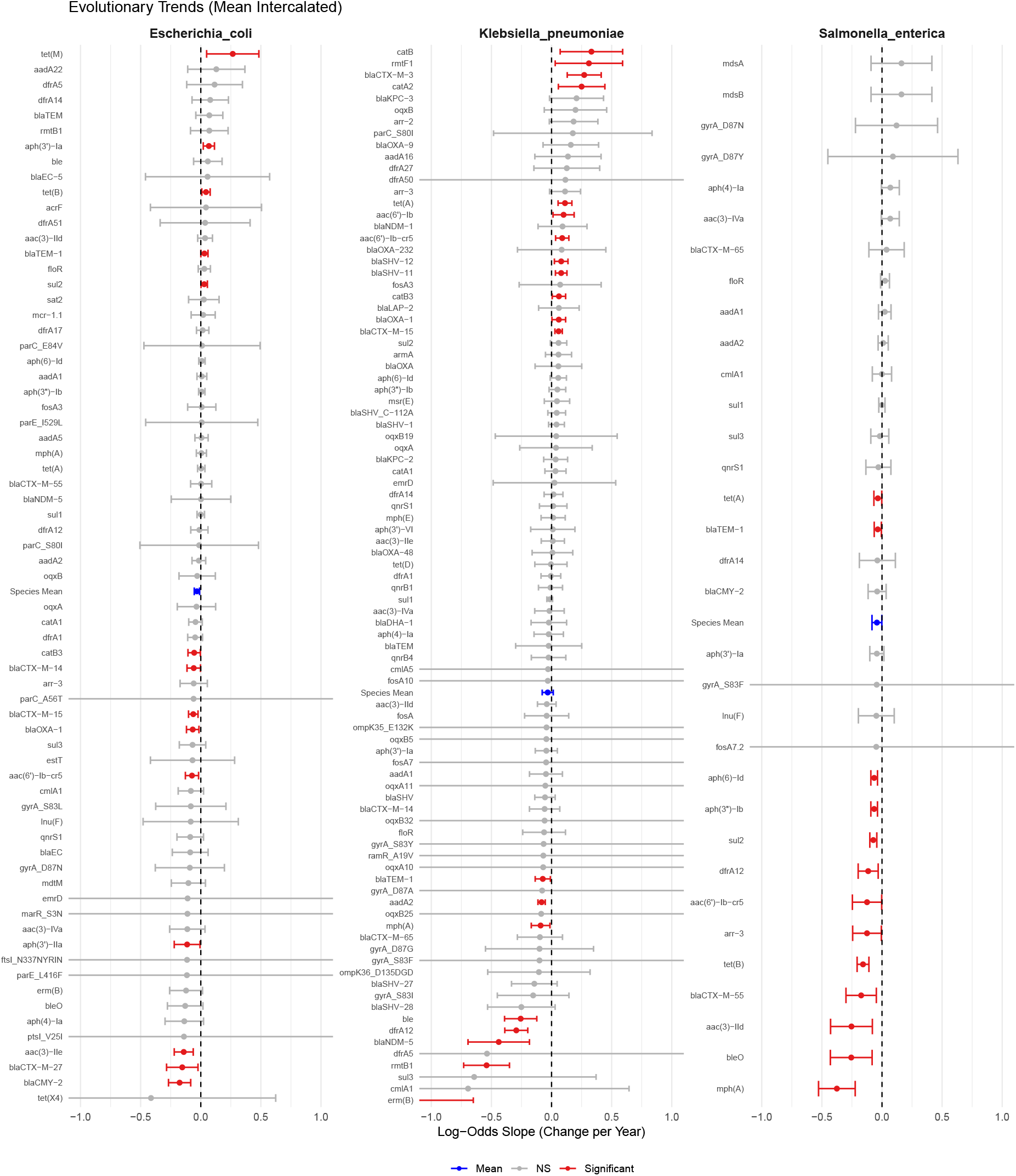
Gene-specific evolutionary trajectories of high-prevalence resistance determinants. Forest plot displaying for each species the temporal shift in genomic location for individual AMR genes with ≥ 100 total observations. Points represent the estimated annual log-odds slope (*β*_1_); error bars indicate 95% Wald confidence intervals. Red markers denote trends significantly different from zero (*P* < 0.05) while grey markers show slopes not significantly different from zero. The blue dot represents the mean slope across all analyzed genes within the species and a 95% confidence interval. Positive values indicate an increasing probability of plasmid carriage over time, while negative values indicate increasing chromosomal association.

## Supplementary Information

### 1 Supplementary Note A: Analytical derivation of the probability of successful plasmid invasion

Here we derive the probability that a single plasmid-carrying cell (*P*), introduced into a resident sensitive population via an exterior conjugation event, successfully establishes a stable lineage. We treat this as a stochastic birth-death process where the invader is initially rare.

#### 1.1 The Resident Sensitive Equilibrium

We assume the population is initially in a stable equilibrium consisting of sensitive (*S*) cells. At this equilibrium, the per capita birth rate must equal the per capita death rate (*d*). Using the logistic growth model, the equilibrium population size *N*_*eq,S*_ is:

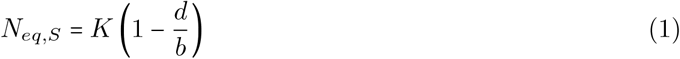

where *b* is the basal birth rate and *K* is the carrying capacity. This resident population defines the competitive environment through the logistic term:

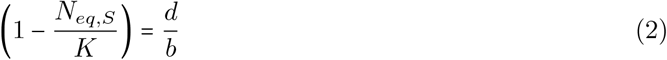

#### 1.2 Invader Kinetics and Probability of Establishment

An invading plasmid-carrier (*P*) has an intrinsic birth rate of *b*_*P*_ *= b(*1+*m* − *c*_*p*_). Its effective birth rate, *b*_*P*,eff_, is scaled by the logistic growth (Eq. 2):

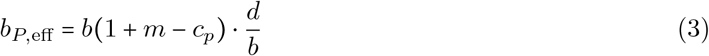

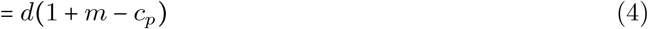

The effective death rate is the constant per capita rate, *d*_*P*,eff_ = *d*.

For a stochastic birth-death process, the probability of establishment *p*_*E,p*_ (the probability of avoiding early stochastic extinction) is given by:

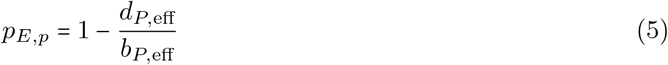

Substituting the effective rates derived above:

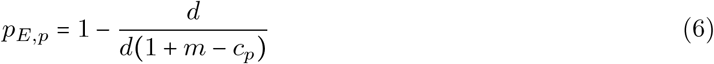

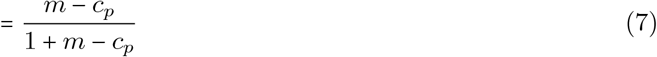

This result shows that the probability of a single plasmid introduction leading to fixation is determined by the ratio of its net selective advantage (*m* − *c*_*p*_) to its relative growth rate. If the metabolic cost of the plasmid exceeds the benefit of resistance (*c*_*p*_ > *m*), the probability of establishment is zero.

### 2 Supplementary Note B: Analytical derivation of the probability of successful transition from plasmid to chromosome resistance

Here, we derive an analytical approximation for the probability of evolutionary rescue, *P*_rescue_. This is the probability that a single plasmid-to-chromosome recombination event (*P* → *B*) ultimately leads to the successful establishment of a chromosomal (*C*) lineage. The derivation follows the three key steps outlined in the main text, using an approximation for the total probability.

#### 2.1 The Resident Equilibrium Environment

The success of the rescue pathway depends on the environment set by the resident population. We assume the system is in a stable equilibrium dominated by plasmid-carriers. Under the assumption of a low plasmid segregation loss rate (*τ*_*p*_ ≪ 1), we can approximate the resident population as being composed almost entirely of plasmid-carrying (*P*) cells. The intrinsic birth rate of this resident population is *b*_*P*_ = *b*(1 + *m* − *c*_*p*_). The total population size at this equilibrium, *N*_*eq,P*_, is therefore:

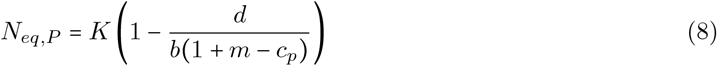

This equilibrium state defines the logistic suppression experienced by any new invader:

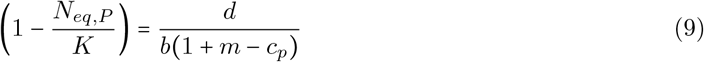

##### Step 1: The Progeny of the Maladapted Intermediate (*E* [*X*])

The rescue lineage is initiated by a single cell carrying both elements (*B*). The expected total progeny size of this subcritical lineage, *E* [*X*], is given by *d*_*B*,eff_ /(*d*_*B*,eff_ − *b*_*B*,eff_).

The intrinsic birth rate of a *B* cell is *b*_*B*_ *= b*(1+*m* − *c*_*p*_ − *c*_*c*_). Its effective birth rate is this value scaled by the logistic suppression from the *P*-dominated environment (Eq. 9):

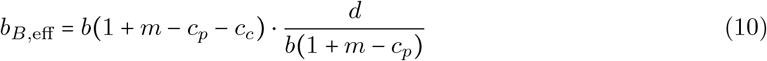

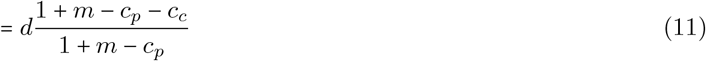

A *B* cell is removed from the *B* lineage by natural death (rate *d*), therefore *d*_*B*,eff_ = *d*. The lineage is subcritical if *d*_*B*,eff_ > *b*_*B*,eff_.

Substituting these effective rates into the formula for the mean progeny size yields the full expression:

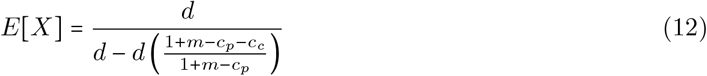

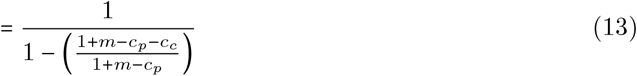

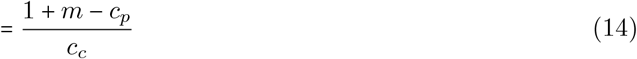

##### Step 2: The Rescue Event (*p*_rescue_)

For any single *B* cell in the progeny, the probability that leads to a single *C* cell is simply the probability of loss of the plasmid upon replication *q*_*L,p*_. Note that if the progeny is of size *x*, this means that there are *x* − 1 events of a *B* cell replicating, as the first one was created by transposition rather than replication.

##### Step 3: Establishment of the Rescued Lineage (*p*_*C*_)

Once a *C* cell is produced, its probability of establishing a successful lineage is *p*_*C*_ *=* 1−*d*_*C*,eff_/*b*_*C*,eff_. Its effective rates are calculated in the same resident *P*-dominated environment.

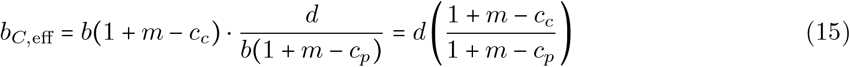

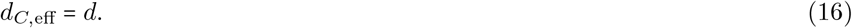

The probability of establishment is therefore:

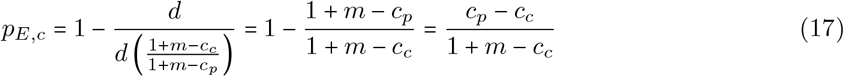

#### 2.2 The Overall Probability of Rescue (*P*_rescue_)

The total probability of the rescue pathway succeeding is the probability that at least one cell in the sub-critical *B* lineage loses the plasmid when created. It is the complementary probability to the probability of all *x* − 1 daughter *B* cells either not losing the plasmid, or losing it and then going extinct. The full probability therefore depends on the distribution of *X*, and gives:

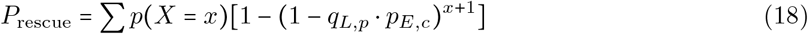

Which can be approximated in the case of a small probability of emergence *q*_*L,p*_ *p*_*E,c*_, using the final expressions for the mean progeny size from Equations (14) - (17), to give the expression:

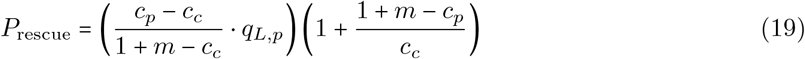

### Supplementary Note C: Sensitivity analysis for chromosome gene parameters

The rate at which a resistance gene transitions from the plasmid to the chromosome depends closely on the parameters governing the chromosomal allele. As shown in Figure S1, increasing the fitness cost associated with the chromosomal element (*c*_*c*_) approximately linearly extends both the period of plasmid dominance and the total time to chromosomal fixation. Analytically, a higher *c*_*c*_ reduces the selective advantage of the rescued *C* lineage over the resident *P* lineage, directly reducing its establishment probability *p*_*E,c*_ (as derived in Eq.(17)). A higher *c*_*c*_ also reduces the progeny size of *B* cells, the necessary intermediate step for chromosome resistance to emerge. Similarly, the transposition rate (*q*_*T*_, denoted as *ϕ* in Fig. S1) acts as the primary mutational bottleneck for this transition. Lower transposition rates decrease the initial production of intermediate *B* cells, thereby increasing the waiting time for a successful integration event and prolonging the transient phase of plasmid dominance.

**Figure S1:**
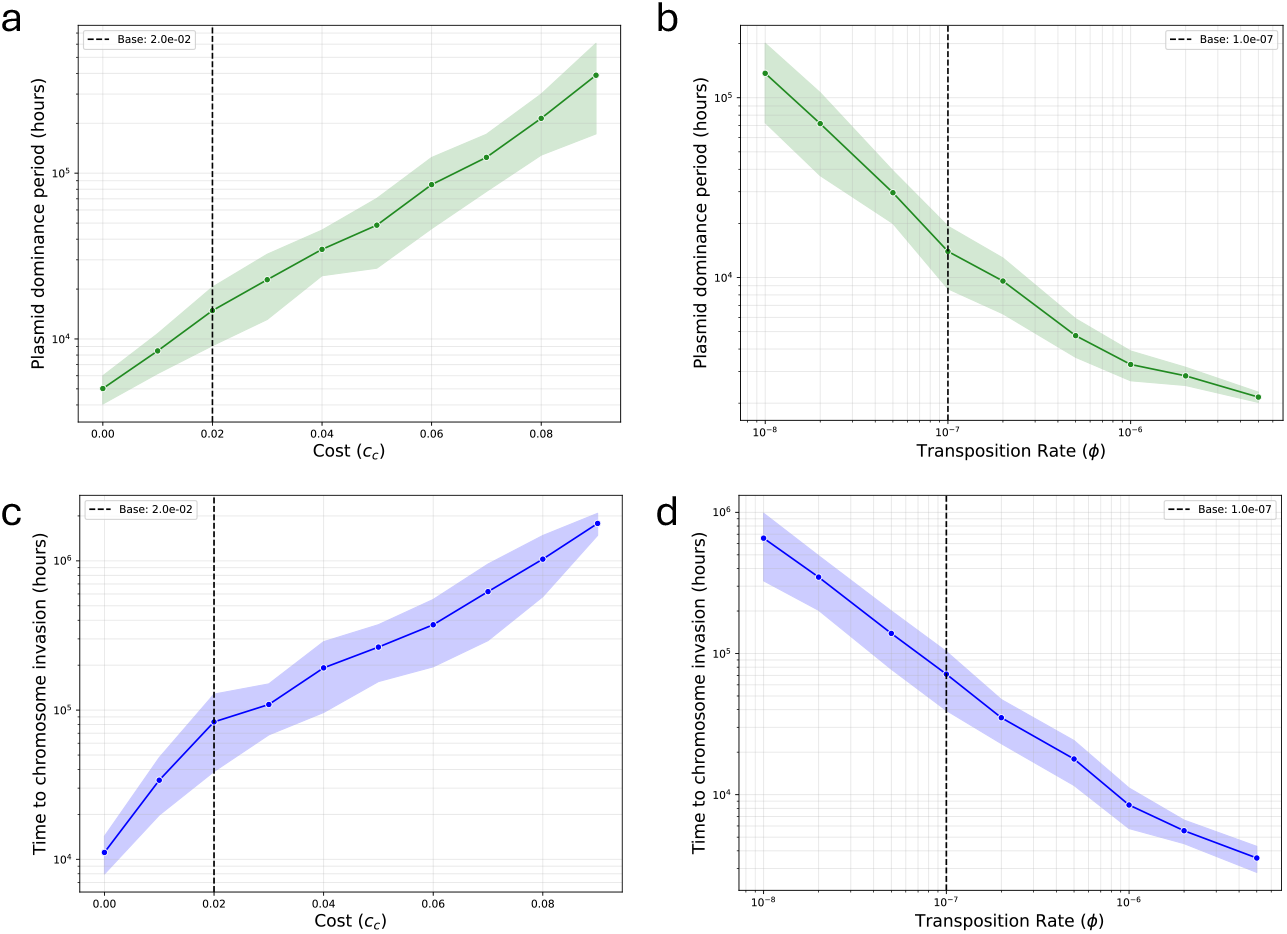
Sensitivity of chromosomal resistance fixation to transposition rate and chromosome-borne gene cost. Impact of varying the chromosome-borne gene cost (*c*_*c*_) (a, c) and the transposition rate (*q*_*t*_, denoted as *ϕ*) (b, d) on the duration of plasmid dominance (a, b) and the total time to chromosome invasion (c, d). Solid lines represent the mean across 50 stochastic simulations, with shaded regions indicating ± SD. Baseline parameter values (*c*_*c*_ *=* 0.02, *ϕ =* 10^−7^) are indicated by vertical dashed lines.

### Supplementary Note D: Considering a strong HGT rate for chromosomal genes

Our base model assumes that horizontal gene transfer for chromosomal genes (*γ*_*c*_) is negligibly low compared to plasmid conjugation (*γ*_*p*_). To verify that this asymmetry is not an artificial driver of our results, we performed a sensitivity analysis by varying the chromosomal HGT rate (Figure S2). The time to chromosomal fixation remains largely insensitive to *γ*_*c*_ until this rate approaches 10^−3^ relative to the plasmid conjugation rate. Given that empirical estimates generally place chromosomal HGT several orders of magnitude lower than plasmid conjugation, this confirms that the slower, vertical-transmission-limited sweep of chromosomal resistance is robust to realistic levels of background chromosomal recombination.

**Figure S2:**
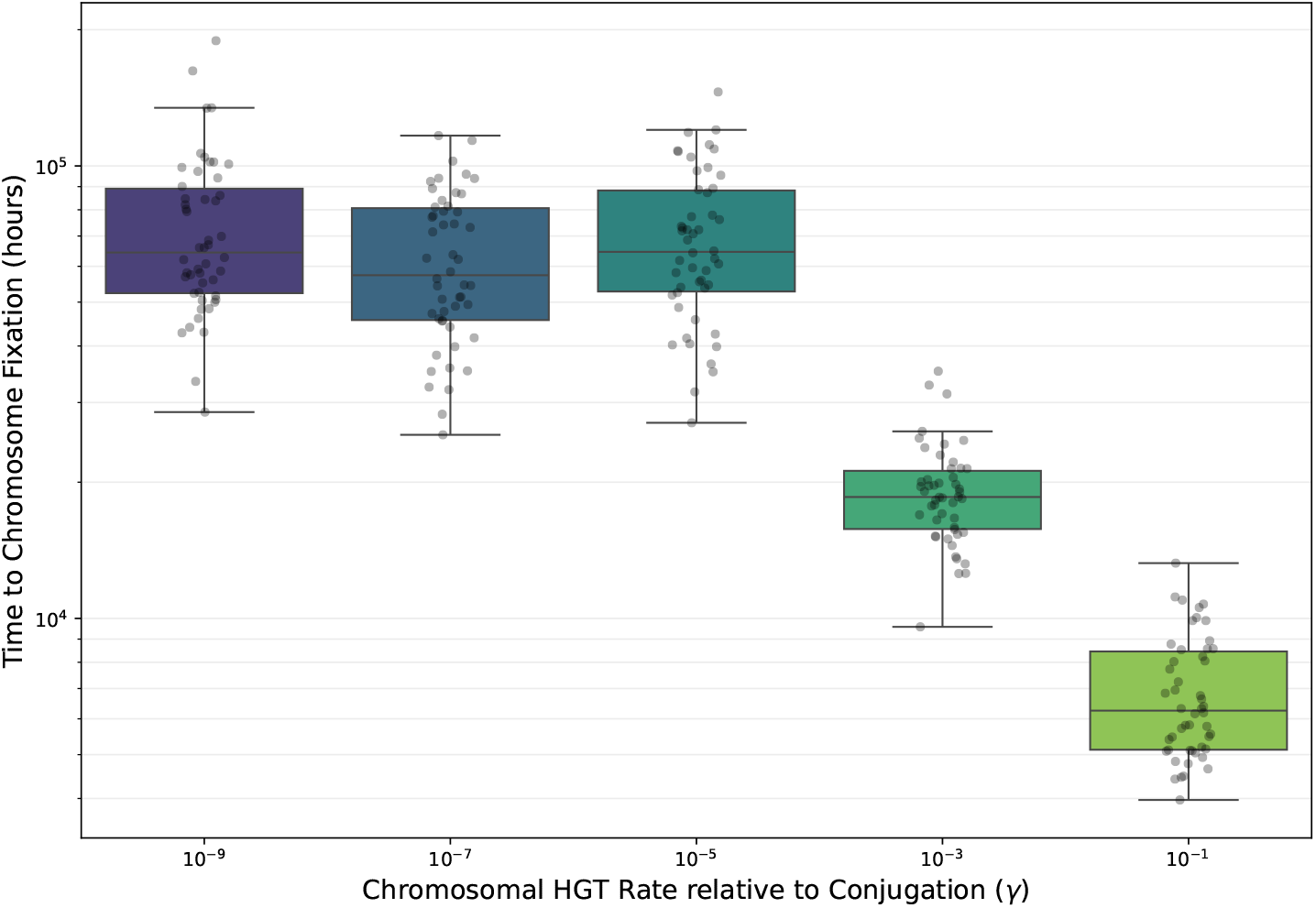
Sensitivity of chromosomal resistance fixation to horizontal gene transfer (HGT) rates. Impact of varying the chromosomal HGT frequency on the time required for chromosomal resistance alleles to reach population-wide fixation (*T*_*C*_). The rate of chromosomal HGT is expressed as a factor relative to the baseline plasmid conjugation rate (*γ =* 10^−12^). Time to fixation is shown on a logarithmic scale. Fixation is defined as the chromosomal resistance allele frequency exceeding the dynamic threshold for *n =* 10 strains (*f*_*C*_ *>* 0.99). Boxplots indicate the median and interquartile range (IQR), with whiskers extending to 1.5× IQR. Individual markers represent independent stochastic simulation replicates (10 per condition).

### Supplementary Note E: Sensitivity analysis for plasmid parameters

We also explored the sensitivity of our results to the intrinsic properties of the plasmid (Figure S3). As expected from our analytical approximations (Eq. (19)), increasing the metabolic cost of the plasmid (*c*_*p*_) strongly penalizes the plasmid-carrying resident population as it increases the relative fitness advantage of a chromosome-resistant *C* cell. This drastically accelerates the transition from plasmid to chromosome dominance. Similarly, an increased segregation loss increases the probability for a *B* cell to create a *C* cell upon replication, also favouring the transition to chromosome-borne resistance. Varying the plasmid conjugation rate (*γ*_*p*_) has very little impact on the timescale of chromosomal fixation. While conjugation is critical for the initial invasion of the plasmid into the naive population, it is not strong enough to revert within-strain chromosome dominance.

**Figure S3:**
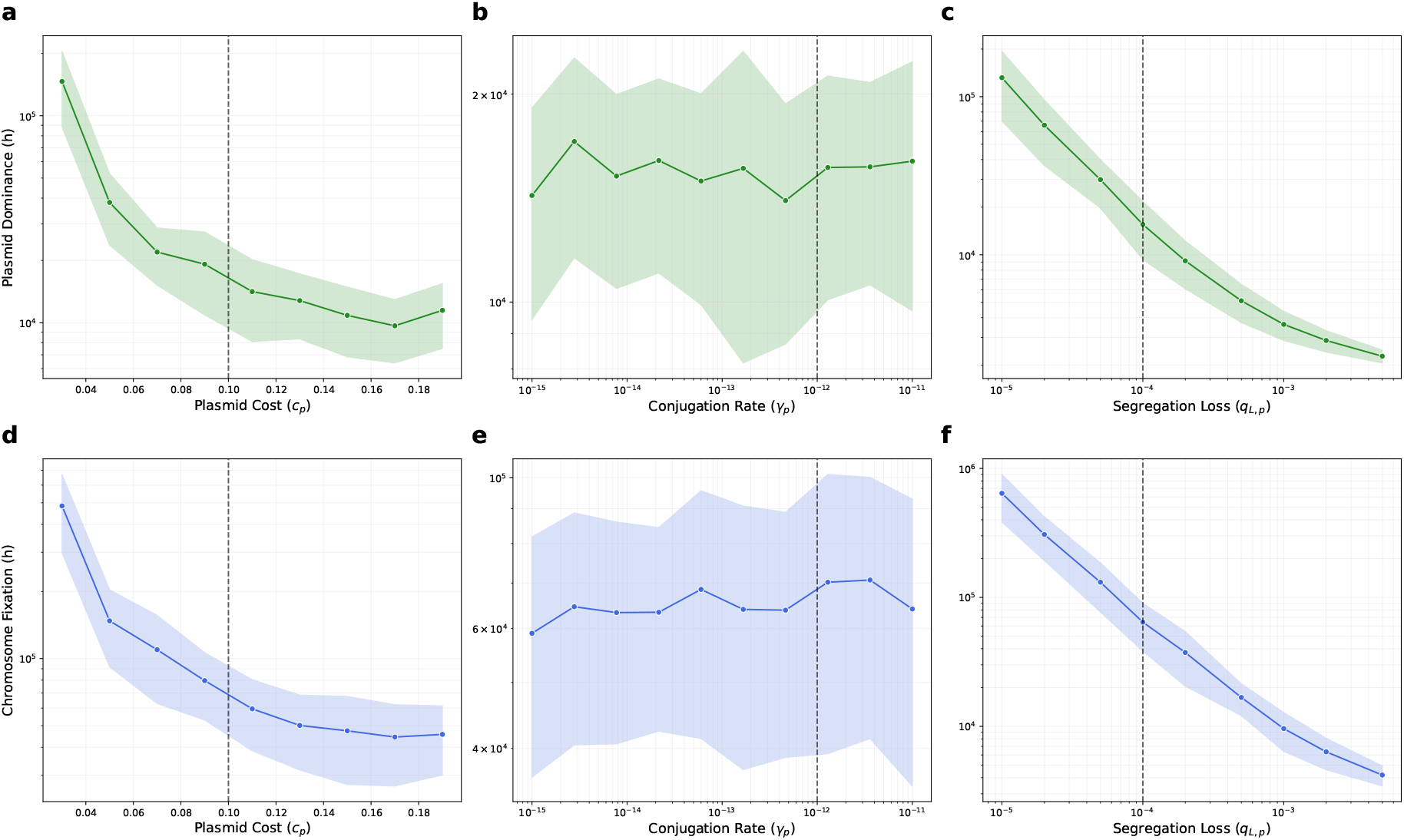
Sensitivity of chromosomal resistance fixation to plasmid cost, conjugation rate, and loss rate. The plots display the plasmid dominance period (a, b, c) and the time to chromosome fixation (d, e, f) as a function of plasmid cost *c*_*p*_ (a, d), conjugation rate *γ*_*p*_ (b, e), and segregation loss rate *q*_*L,p*_ (c, f). Solid lines represent mean values across simulations, with shaded regions indicating ± SD. Vertical dashed lines mark the baseline parameter values used in the main text.

### Supplementary Note F: Considering variation in population size

Because the evolutionary rescue of the chromosomal allele relies on rare stochastic events (transposition and subsequent lineage establishment), the dynamics are highly sensitive to the absolute population size. We varied the system’s carrying capacity (*K*) to examine this demographic effect (Figure S4). As the carrying capacity increases, the absolute number of plasmid-carrying cells at equilibrium increases, which proportionally increases the absolute number of transposition events occurring per unit time in the population. Consequently, the waiting time for the emergence of a successful chromosomal lineage scales inversely with the carrying capacity. Despite these varied timescales, the qualitative trajectory of the plasmid-to-chromosome transition remains consistent across all tested population sizes.

**Figure S4:**
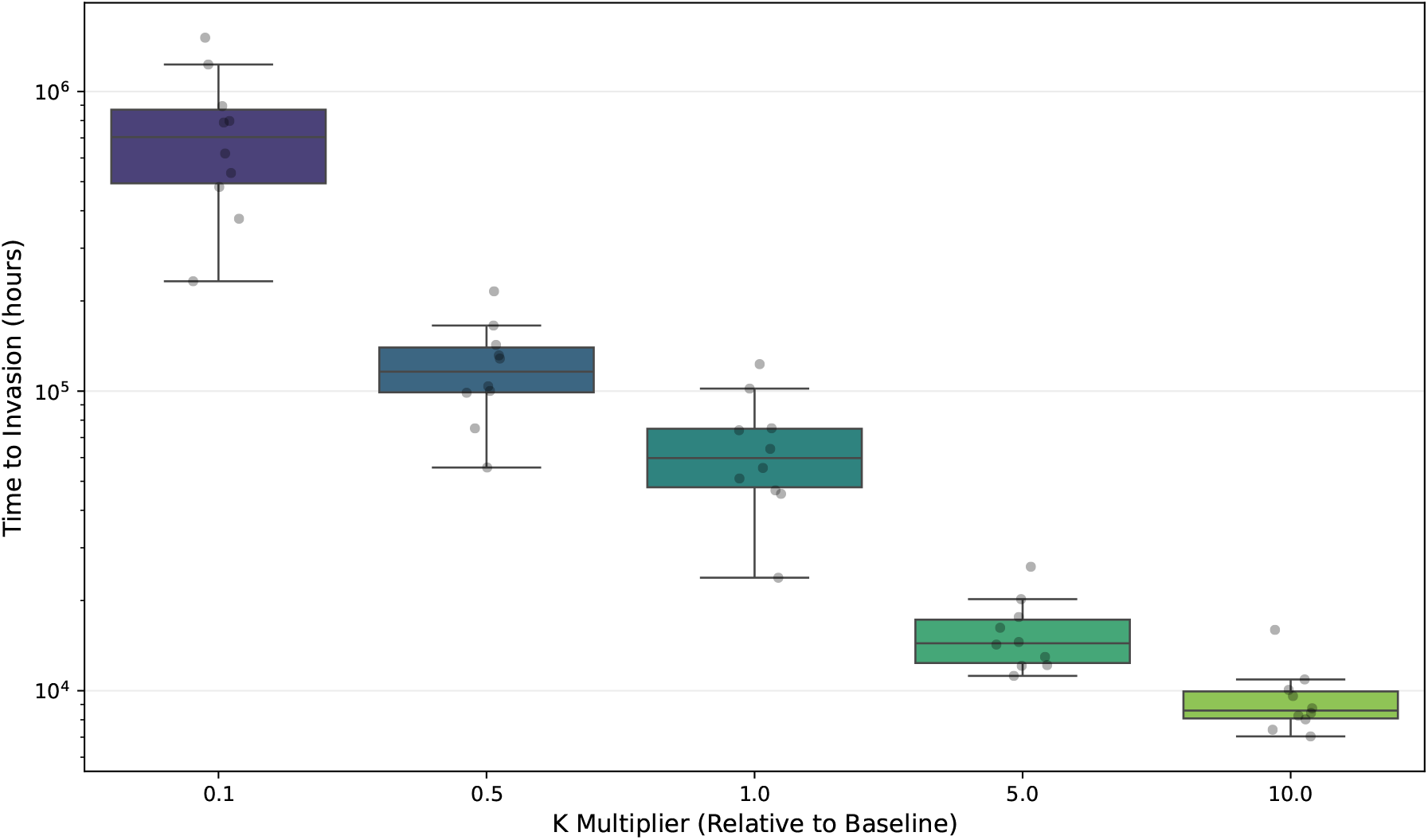
Gene location dynamics and population size. Relationship between the carrying capacity (*K*) and the time required for chromosomal resistance alleles to reach population fixation (using a frequency threshold of 0.99). The baseline value used is *K =* 10^8^cells.ml^−1^ This threshold ensures that all strains have been invaded and accounts for a portion of sensitive cells still remaining at equilibrium. Boxplots represent the median and interquartile range (IQR), with whiskers extending to 1.5× IQR. Overlaid circular markers indicate the results of individual stochastic simulation replicates (50 per condition). Simulations were initialized with a sensitive population consisting of 10 strains, and a 10^6^ inoculum of plasmid-bearing cells in a single strain. Fixation times are expressed in simulation time units (hours).

### Supplementary Note G: Considering multi-drug resistance (MDR)

In this version of the model, we assume the existence of another plasmid carrying two resistances, *A* and *B*. Both genes behave the same, and the same as the single resistance does in the rest of this work. A cell can harbor only one plasmid at a time, and a pre-existing plasmid prevents the acquisition of another via conjugation.

The state of a cell is defined by (*p, c*) where *p, c* ∈ {0, 1, 2}, representing none, Gene *A*, or Both (MDR).

- **Sensitive**: (0, 0); *w =* 1
- **Plasmid A**: (1, 0); *w =* 1 + *m* − *c*_*p*_
- **Chromosome A**: (0, 1); *w* = 1 + *m* − *c*_*c*_
- **Plasmid A + Chrom A**: (1, 1); *w =* 1 + *m* − (*c*_*p*_ *+ c*_*c*_)
- **MDR Plasmid**: (2, 0); *w =* 1 + 2*m* − (*c*_*p*_ *+ c*_*c*_)
- **MDR Plasmid + Chrom A**: (2, 1); *w =* 1 + 2*m* − (*c*_*p*_ *+* 2*c*_*c*_)
- **Chromosome AB**: (0, 2); *w =* 1 + 2*m* − 2*c*_*c*_
- **MDR Plasmid + Chrom AB** : (2, 2); *w =* 1 + 2*m* − (*c*_*p*_ *+* 3*c*_*c*_)
- **Plasmid A + Chrom AB** : (1, 2); *w =* 1 + 2*m* − (*c*_*p*_ *+* 3*c*_*c*_)

The dynamics for strain *i* are:

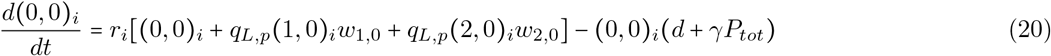

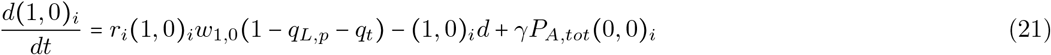

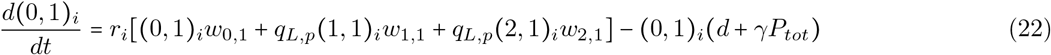

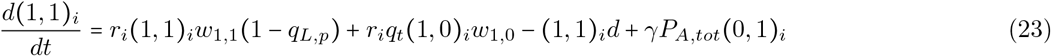

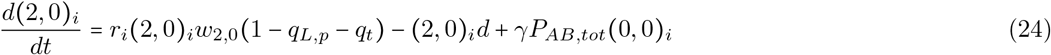

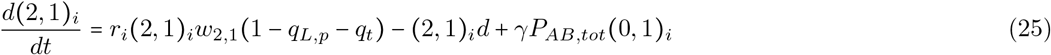

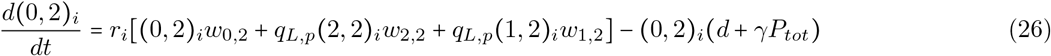

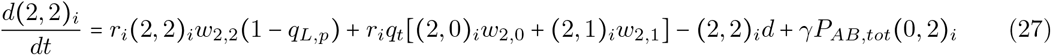

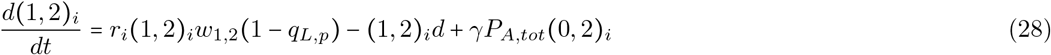

with the donor pools

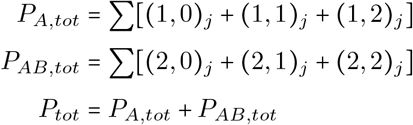

We initialize the simulations with 10^6^ (1, 0)_1_ cells, similarly to the main model. When the frequency of resistance *A* on the chromosome becomes higher than 0.99, an inoculum of 10^6^(2, 1)_1_ cells is made, to mimic the introduction of a new multi-resistant plasmid in that strain.

#### Biological Assumptions

##### Strict Incompatibility

No plasmid displacement is permitted. Conjugation only occurs if the recipient cell has a plasmid mask of 0. Note that interactions between the two plasmids and whether we allow for co-colonisation, displacement or (as is the case) complete incompatibility has very little impact on the dynamics, as we introduce the *AB* plasmid when the *A* plasmid is going to, and close, to extinction.

##### Linked Transposition

Recombination *q*_*t*_ is replication-coupled. For the MDR plasmid (index 2), a single event copies both determinants to the chromosome.

#### No between-cell chromosome recombination, no chromosomal gene loss

The estimated rates for between-cell recombination for a chromosomal gene are so small that we assume this process is negligible in this model. The loss of a chromosomal gene, although more frequent in reality, has a negligible effect on the dynamics (they can be compared to back mutations, deleterious) and is also not considered here.

### Supplementary Note H: Stochastic Modeling and Numerical Implementation

#### Finite Population Framework

To account for demographic stochasticity and the probabilistic nature of rare evolutionary transitions, we implement the model as a Continuous Time Markov Chain (CTMC) with discrete states. The state of the system is defined by the vector **N**(*t*), containing the discrete number of cells for each of the four genotypes within each strain *i*: sensitive cells (*S*_*i*_), plasmid-carriers (*P*_*i*_), chromosomal-carriers (*C*_*i*_), and double-carriers (*B*_*i*_).

The total population size 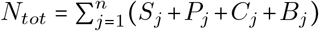 is finite and varies stochastically around the carrying capacity *K*. The per capita birth rate of each genotype is regulated by a logistic constraint and strain-level negative frequency-dependent selection (NFDS). For a strain *i*, we define the growth factor *r*_*i*_ as:

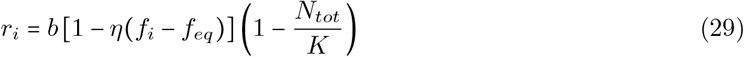

where *b* is the basal birth rate, *f*_*i*_ is the frequency of strain *i*, and *η* is the strength of NFDS.

#### Event Propensities and Inheritance Logic

We categorize stochastic transitions into rate-based events (Poisson processes) and replication-coupled events (conditional Binomial processes).

##### Rate-based transitions

Death and horizontal gene transfer (HGT) occur at fixed rates proportional to the density of donors and recipients. The propensity ℛ for these events is:

- **Death:** ℛ_*death*_ = *dN*_*G,i*_ for any genotype *G* of strain *i*.
- **Conjugation:** ℛ_*conj*_ = *γ*_*p*_*P*_*tot*_*N*_*rec,i*_ where the recipient *N*_*rec,i*_ ∈ {*S*_*i*_, *C*_*i*_}.
- **Chromosomal HGT:** ℛ_*hgt*_*c*_ = *γ*_*c*_*C*_*tot*_*N*_*rec,i*_ where the recipient *N*_*rec,i*_ ∈ {*S*_*i*_, *P*_*i*_}.

##### Replication-coupled transitions

Genetic transitions involving plasmid loss, chromosomal loss, and transposition are modeled as probabilities per replication event. For a genotype *G* with relative fitness *w*_*G*_, the total number of division events *κ*_*birth*_ occurring in a time interval *τ* is drawn from a Poisson distribution:

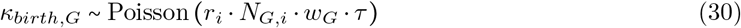

The genetic identity of the resulting offspring is determined by a sequence of Binomial draws, ensuring that a single birth event results in exactly one daughter cell. For a plasmid-only carrier *P*_*i*_ (*w*_*P*_ = 1 + *m* − *c*_*p*_), the offspring are distributed as follows:

1. **Plasmid loss:** *κ*_1_ ~ Binomial(*κ*_*birth*_, *q*_*L,p*_) daughter cells revert to *S*_*i*_.
2. **Transposition:** *κ*_2_ ~ Binomial(*κ*_*birth*_ − *κ*_1_, *q*_*t*_) daughter cells become *B*_*i*_.
3. **Successful inheritance:** *κ*_*birth*_ − *κ*_1_ − *κ*_2_ daughter cells remain *P*_*i*_.

#### Numerical Implementation (*τ*-leap)

Trajectories are simulated using the *τ*-leap approximation of the Gillespie algorithm. Propensity functions are assumed constant over a small interval *τ =* 0.5 hours. The update procedure at each step is:

i. Calculate event rates for all rate-based transitions (Table S1).
ii. Draw the number of occurrences for each event *e* from *κ*_*e*_ ∼ Poisson ~ (ℛ_*e*_*τ*).
iii. For birth events, execute the conditional Binomial inheritance logic described in the “Replication-coupled transitions” section above.
iv. Update the population vector: **N**(*t* + *τ*) = **N**(*t*) + ∑ *κ*_*e*_**v**_*e*_, where **v**_*e*_ is the state change vector.

**Table S1:** Stochastic Jumps and Rates. Summary of the transitions in the CTMC. *w*_*G*_ represents the relative fitness of genotype *G*. For replication-coupled events, the rate shown is the effective mean rate derived from the Poisson-Binomial process. *q*_changes_ denotes all the probabilities upon replication that lead to a daughter cell different from its parent.

| Event | Jump ($dN_i$) | Effective Rate |
| --- | --- | --- |
| *Replication-coupled* |  |  |
| Successful Birth | $N_{(p,c),i} \rightarrow +1$ | $r_i N_{(p,c),i} w_{p,c} (1 - \sum q_{\text{changes}})$ |
| Plasmid Loss | $N_{(0,c),i} \rightarrow +1$ | $r_i N_{(p,c),i} w_{p,c} q_{L,p}$ |
| Chromosomal Loss | $N_{(p,0),i} \rightarrow +1$ | $r_i N_{(p,c),i} w_{p,c} q_{L,c}$ |
| Transposition | $N_{B,i} \rightarrow +1$ | $r_i (N_{P,i} w_P + N_{C,i} w_C) q_t$ |
| *Rate-based* |  |  |
| Death | $N_{G,i} \rightarrow -1$ | $dN_{G,i}$ |
| Conjugation | $N_{rec,i} \rightarrow -1, N_{plas,i} \rightarrow +1$ | $\gamma_p (P_{tot}/K) N_{rec,i}$ |
| Chromosomal HGT | $N_{rec,i} \rightarrow -1, N_{chrom,i} \rightarrow +1$ | $\gamma_c (C_{tot}/K) N_{rec,i}$ |

